# Landrace and bred accessions of allotetraploid sour cherry (*Prunus cerasus* L.) reveal variation in subgenome dosage and subgenome expression bias

**DOI:** 10.64898/2026.02.19.706907

**Authors:** Kathleen E.B. Rhoades, Charity Z. Goeckeritz, Kevin A. Bird, Alan Yocca, Patrick Edger, Amy Iezzoni

**Author notes:** Savanna Institute, Madison, WI 53704. Address all correspondence to: Amy Iezzoni Department of Horticulture Michigan State University 1066 Bogue St A288 East Lansing, MI 48824.

## Abstract

Subgenome dominance is a phenomenon observed in many allopolyploids where one parental genome exhibits stronger influence over phenotype than the other parental genomes. This may present as preferential retention of one subgenome through fractionation, replacement via homoeologous exchange, or as subgenome expression bias, where one subgenome is expressed at a higher abundance compared to other subgenomes. Sour cherry (*Prunus cerasus*) is an allotetraploid fruit tree species resulting from an interspecific cross between extant relatives of ground cherry (*P. fruticosa*) and sweet cherry (*P. avium)*. Prior comparative genomic analyses suggest that the sour cherry cultivar Montmorency contains three subgenomes. Subgenomes A and A’, each present in one copy, are derived from a *P. fruticosa-*like ancestor, and B, present in two copies, is derived from a *P. avium*-like ancestor. In this study we investigated the subgenome dynamics of the three subgenomes of sour cherry in four diverse landraces and two cultivars, including ‘Montmorency’. We found evidence of 26 homoeologous exchange events and five whole-homoeolog replacements relative to ‘Montmorency’ in three of the six accessions. We also detected subgenome expression bias favoring the A and A’ subgenomes over the B subgenome, the magnitude of which differs between accessions and changes over the course of fruit development. Lastly, we show differences in dosage variation and expression bias of four previously-described genes in ‘Montmorency’ associated with fruit softening, a key trait in this crop. These findings on subgenome dominance offer valuable insights into how this phenomenon may influence traits important for sour cherry breeding.

## Introduction

Allopolyploidy, characterized by the presence of three or more complete sets of chromosomes derived from two or more diploid progenitor species, is a common phenomenon in plants [1–5]. The success of allopolyploidy is generally attributed to the novel phenotypic variation that emerges from the interactions between two or more diverse subgenomes that evolved separately, as well as the subsequent evolution of the newly formed polyploid genome [6]. Allopolyploid organisms may exhibit a higher competitive ability or expanded geographical range compared to their diploid progenitors [7–10]. However, newly-formed allopolyploids (i.e. neopolyploids) are frequently observed to be genetically unstable, owing largely to the challenge of re-establishing stable meiosis after an allopolyploidy event [11]. Homoeologous chromosomes (i.e. homologs derived from different progenitors) may pair during meiosis and recombine, resulting in a homoeologous exchange (HE) event. Selection may act upon the results of HE, leading to diverse evolutionary trajectories. These range from complete homogenization of the ancestral subgenomes to preferential retention of one subgenome over others [11–13]. Furthermore, neopolyploid formation can induce not only structural changes due to HE, but also result in alterations in DNA methylation patterns and transposable element activity. These modifications can further contribute to deviations from progenitor gene expression patterns [14,15].

Gene expression levels may not be equal between subgenomes in allopolyploids, a phenomenon referred to as subgenome expression bias [16]. A bias in homoeolog expression can contribute to the preferential retention of one subgenome. One model suggests that selection may act against genes with lower expression levels, as these are more likely to be lost with minimal fitness consequences [17]. Over time, allopolyploid genomes tend to reach a more stable, diploid-like state through selection favoring meiotic configurations that generate balanced gametes and therefore ensure sufficient fertility [18]. The vast majority of allopolyploid seed and fruit crops have been selected for mechanisms that promote stable chromosome pairing during meiosis, typically through bivalent pairing between homologous chromosomes. This results in high fertility, which is vital for production, as fertilization is generally essential for fruit/grain set and development.

Sour cherry (*Prunus cerasus* L., 2*n* = 4𝑥 = 32) is an allopolyploid perennial fruit crop with well-documented meiotic irregularities [19–22]. Sour cherry was formed from interspecific hybridization between an ancestor of the extant diploid sweet cherry (*P. avium* L., 2*n* = 2𝑥 = 16) and an ancestor of the extant allotetraploid ground cherry (*P. fruticosa* Pall., 2*n* = 4𝑥 = 32) where their native distributions overlap in Eastern Europe and Western Asia [23,24]. Chloroplast data suggests this hybridization event occurred at least twice, and that *P. fruticosa* was most commonly the maternal parent [25,26], leading to a hypothesis that the hybridization occurred when a *P. avium* individual failed to properly reduce its gametes and produced a 2*n* pollen, which fertilized the *n* ovum of *P. fruticosa*.

The low fertility and poor fruit set exhibited by sour cherry have been attributed to low gamete viability as a result of meiotic irregularities. Studies of meiosis in sour cherry show varying rates of univalents and multivalent chromosome pairing, resulting in aneuploid gametes [19,20,22,27]. Sour cherry exhibits disomic and tetrasomic patterns of inheritance, further supporting occasional pairing between homoeologs at meiosis [28]. The consequence of the low maternal gamete fertility in cherry is particularly severe as cherry is not a multi-seeded fruit such as tomato or blueberry. Instead, cherry and all *Prunus* species have just two ovules, and typically one degenerates, leaving one seed inside the endocarp. Fruit set requires successful fertilization and the initiation of embryo development. Therefore, any disruption that results in a non-viable ovule or zygote will result in the flower failing to develop into a fruit.

All commercial sour cherry cultivars grown in the major production areas are either landrace cultivars that are hundreds of years old, or a first-generation offspring of a landrace [29]. For example, the sour cherry cultivar ‘Montmorency’, that represents over 99% of sour cherry production in the U.S., is believed to have originated ∼ 600 years ago in France [30]. These landraces, selected by local populations over centuries, were vegetatively propagated and therefore are considered clonal groups. Historically, humans made selections within these clonal groups, as these landrace cultivars tend to have higher fruit set compared to the low fruit set exhibited by sexual offspring. However, unlike the vast majority of allopolyploid crop plants, both natural and human selection have failed to result in cytological diploidization and an increase in fertility. Therefore, the biggest breeding challenge for sour cherry continues to be breeding for high yield, as sour cherry hybrid offspring typically exhibit just ∼2% fruit set while at least 25% fruit set is needed for a commercial crop [31].

The first reference genome for sour cherry (cv. Montmorency) [32] was recently released, providing the opportunity to compare subgenome dynamics in diverse sour cherry accessions. In this work, it was proposed that *P. fruticosa* is itself an allotetraploid that passed one copy of each of its subgenomes to sour cherry. As a result, the ‘Montmorency’ reference genome consists of three sets of eight chromosomes representing the two subgenomes inherited from the *P. fruticosa* progenitor (denoted as A and A’) and the one subgenome inherited from the *P. avium* progenitor (denoted as B)[32]. This finding raises the question of whether the tri-subgenome structure in sour cherry may underlie the observed meiotic irregularities, as it is conceivable that the A and A’ subgenomes would not have paired in the nucleus of the *P. fruticosa* ancestor. Herein, we utilized the ‘Montmorency’ reference genome as a foundation to examine subgenome dynamics, both structural and expression-based, of ‘Montmorency’ and five other sour cherry accessions. We then queried our findings utilizing two sets of genes previously identified in sour cherry. The first set were the well characterized sour cherry self-incompatibility alleles located at the *S*-locus that control the specificity of self-pollen rejection, and ultimately whether a sour cherry selection is self-compatible or self-incompatible [33–35]. The second set of genes were four previously identified expansin genes that were expressed in the final stages of ‘Montmorency’ fruit ripening, suggesting that they may be associated with fruit softening [36]. Expansins are cell wall-modifying enzymes [37,38] that have been associated with fruit ripening and softening processes in tomato [39], pear [40], strawberry [41–43], and peach [44,45], among others. Taken together, we report (*i*) genome-wide subgenome dosage variation among four diverse landrace and two bred sour cherry accessions, (*ii*) patterns of subgenome expression bias in five tissues from those same accessions, and (*iii*) differences in dosage variation and expression bias during fruit development and ripening for four previously characterized expansin genes [36].

## Results

### Predicting chromosome replacements and sites of homoeologous exchange

When short reads were aligned back to the ‘Montmorency’ reference genome, differences in chromosome dosage (copy number) and sites of putative homoeologous exchange (HE) were readily identified in three of the six accessions based on reciprocal changes in read coverage (Fig. 1, Supplementary Fig. S2, Supplementary Table S1). Fig. 2 shows stylized karyotypes of the chromosome dosage and predicted HEs for each accession. As expected, the chromosome subgenome dosage for ‘Montmorency’ is consistently 1A:1A’:2B, reflecting an equal genome contribution from each progenitor species (i.e., *P. fruticosa* and *P. avium*). The sharp spikes and drops in chromosome coverage that were not mirrored in another homoeolog likely represent differences in repeat content between the accession and the reference assembly and were therefore not considered changes in homoeolog dosage. ‘Balaton’ and ‘Oblačinska’ also exhibit a reference-level dosage of all homoeologs, with no predicted chromosome replacements or HEs (Supplementary Fig. S2).

**Figure 1:**
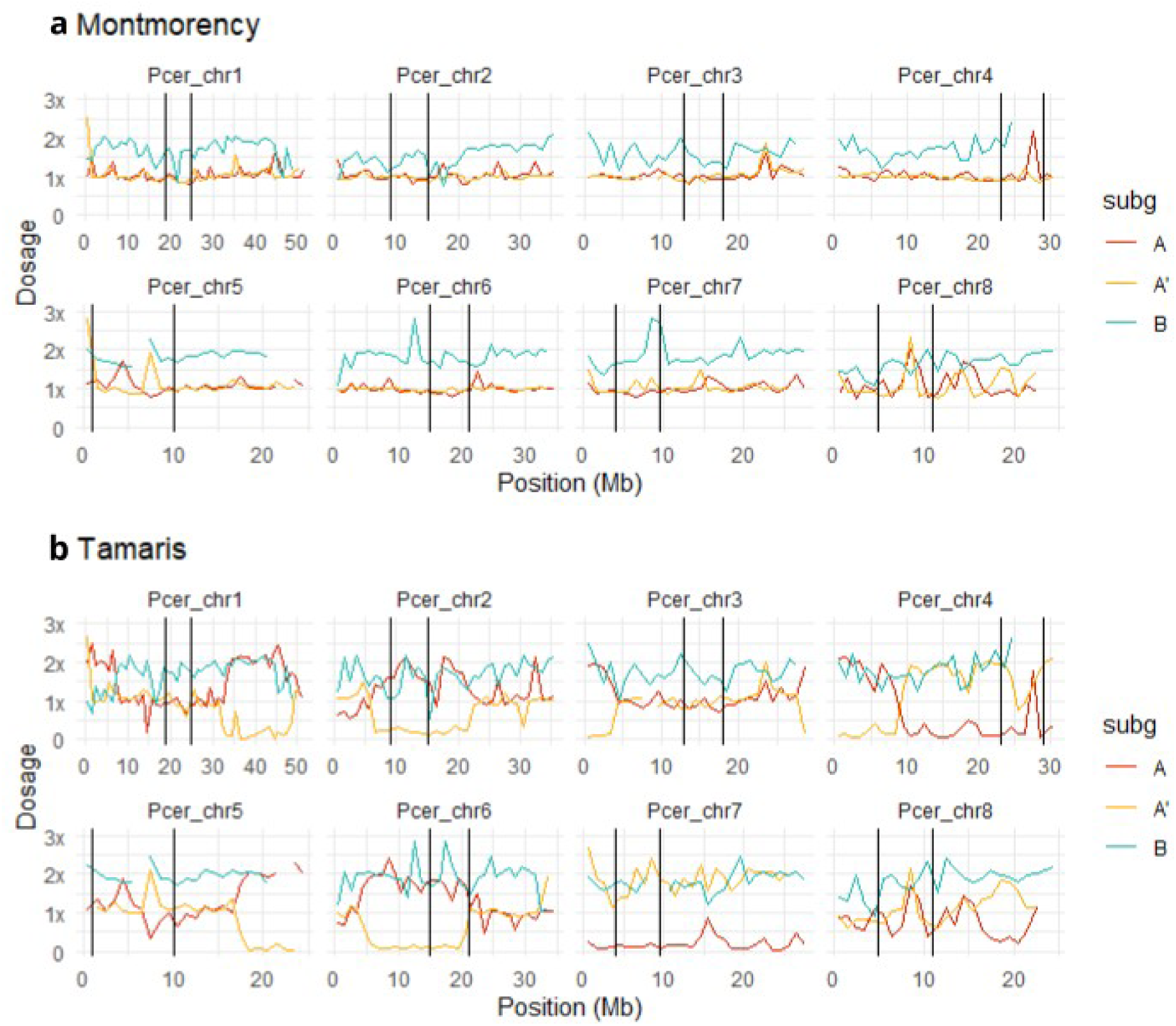
DNA sequence coverage graphs of the eight chromosomes for ‘Montmorency’ and ‘Tamaris’, generated by aligning 2×150bp short reads to the ‘Montmorency’ reference and averaging coverage over 1Mb windows. The 1×, 2×, and 3× dosage lines on the y axis correspond to 20×, 40× and 60× coverage, respectively. The predicted boundaries of the centromeric regions are denoted with black lines and were taken from Goeckeritz et al. (2023). Red, yellow and blue lines denote A, A’ and B homoeolog coverage, respectively. Gaps in the coverage graphs are due to spikes in coverage that peak above the y-axis limit for the plot. The coverage graphs for the other four accessions are in Supplementary Fig. S2. **Alt text:** Two sets of line graphs, one for ‘Montmorency’ and one for ‘Tamaris’. For each set there is one line graph per linkage group and different-colored lines showing the sequence coverage for each subgenome in that linkage group.

**Figure 2:**
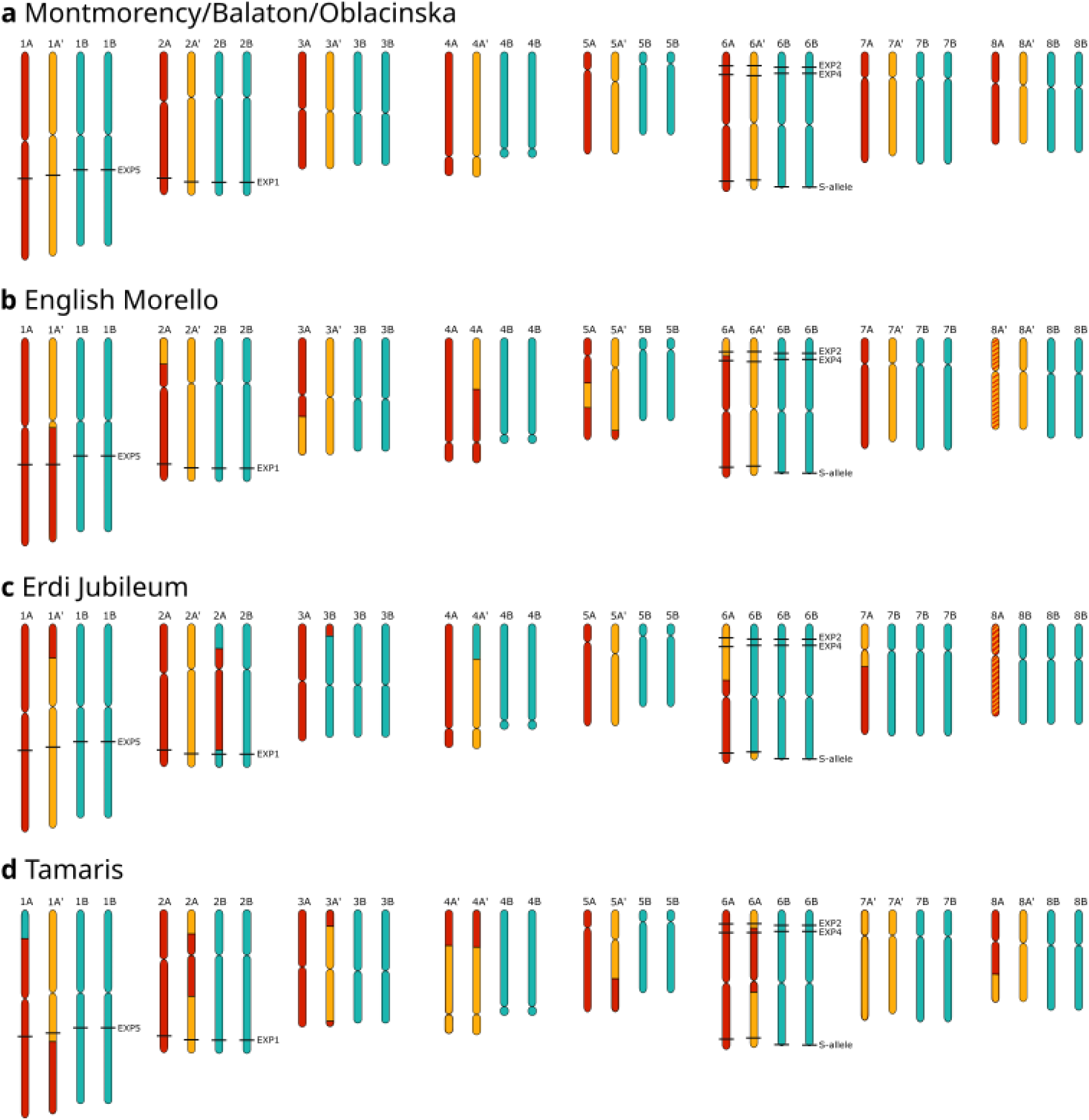
Karyotype illustration of subgenome content and homoeologous exchange sites for each accession based on the coverage shown in Fig. 1 and Supplementary Fig. S2. The chromosome locations of the four expansins first characterized by Yoo et al. (2003) are marked, as well as the location of the *S*-locus. The A, A’ and B homoeologs are colored red, yellow and blue, respectively. Chromosome labels are based on the subgenome identity of the centromeric region. Homoeologs with ambiguous identity (‘English Morello’ chr. 8 and ‘Érdi Jubileum’ chr. 8) are given striped coloring of the two subgenomes contributing to their makeup. **Alt text:** Four depictions of the eight chromosomes of sour cherry, each with four copies colored according to their subgenome identity, labeled a-d. The locations of four expansin genes are marked on chromosomes 1, 2, and 6. Panels b-d, representing ‘English Morello’, ‘Erdi Jubileum’, and ‘Tamaris’, respectively, show crossover events (changes in color) along several chromosomes.

In contrast, a total of 26 HE events and four whole-chromosome replacements were identified in the three remaining accessions (‘Tamaris,’ ‘English Morello,’ and ‘Érdi Jubileum’). Twenty of the 26 HEs detected were between the two *P. fruticosa*-derived subgenomes A and A’, while the other six HEs were between the *P. avium*-derived B subgenome and either the A or A’ subgenomes (Supplementary Table S1a). For example, the centromeric region of ‘Tamaris’ chromosome 1 exhibits the reference-level 1A:1A’:2B subgenome dosage; however, predicted HE events on both chromosome arms result in non-reference subgenome dosage at the ends of each arm (Figs. 1 and 2). Specifically, the predicted HE at the bottom of ‘Tamaris’ chromosome 1 between the A and A’ subgenomes results in a subgenome dosage of 2A:0A’:2B, while the predicted HE at the top of the ‘Tamaris’ chromosome 1 between the A and B subgenomes results in a subgenome dosage of 2A:1A’:1B. All of the HEs identified appear to be unique events, with the exception of exchanges between chromosomes 6A and 6A’ and chromosomes 2A and 2A’ that are common to both ‘English Morello’ and ‘Tamaris’ (Supplementary Table S1b).

In ‘English Morello’ and ‘Érdi Jubileum’, we were not able to make confident dosage predictions for chromosomes 8A and 8A’, as these chromosomes show ambiguous sequence coverage that made assigning homoeolog dosage difficult (Supplementary Fig. S2c, d). ‘Érdi Jubileum’ short read coverage indicates a ∼3× dosage of chromosome 8B and a ∼0.5× dosage each of chromosomes 8A and 8A’. ‘English Morello’ short read coverage indicates a ∼2× dosage of chromosome 8B, a ∼1.5× dosage of chromosome 8A’, and a ∼0.5× dosage of chromosome 8A. Goeckeritz et al. (2023) reported some difficulty in phasing chromosomes 8A and 8A’ due to ambiguity of the Hi-C matrix that may be due to sequence similarity between these two *P. fruticosa*-derived subgenomes.

‘Érdi Jubileum’ and ‘Tamaris’, the two bred cultivars, were the only two accessions to exhibit whole-chromosome replacements (Fig. 1, Supplementary Fig. S2, Fig. 2). ‘Tamaris’ had two copies each of homoeologs 7A’ and 7B and no copies of homoeolog 7A resulting in a subgenome dosage of 0A:2A’:2B for this chromosome. ‘Érdi Jubileum’ exhibited chromosome replacements for five of its eight chromosomes. For chromosomes 3, 6, 7 and 8, three copies of the *P. avium*-derived B homoeologs were present and one homoeolog was either A or A’ from the *P. fruticosa* progenitor. However, for chromosome 2, ‘Érdi Jubileum’ had only one 2B homoeolog, as the other B chromosome was replaced by a second A homoeolog resulting in a chromosome 2 dosage of 2A:1A’:1B. Three of the five chromosome replacements identified in ‘Érdi Jubileum’ were accompanied by HEs on the chromosome arms. For example, the top of one of the three 3B homoeologs has a 3A segment that would have resulted from a HE between these two homoeologs. Despite the high number of HE events and chromosome replacements in ‘Érdi Jubileum’, overall subgenome dosage was still 26% A, 16% A’ and 58% B (Supplementary Table S2).

As an internal control, the previously-characterized *S*-alleles allowed us to further verify our A/A’ versus B subgenome dosage results for the *S*-locus region at the bottom of chromosome 6. *S*-alleles common to both sweet and sour cherry are presumed to have been passed from sweet cherry to sour cherry, and *S*-alleles present in sour cherry but not in sweet cherry are presumed to have been derived from the *P. fruticosa* ancestor. The homoeolog dosage we determined for the *S*-locus region of chromosome 6 for each accession was consistent with the progenitor origin of previously reported *S*-alleles. Specifically, for all five accessions except ‘Érdi Jubileum’ the homoeolog dosage of the *S*-locus region was 1A:1A’:2B which was consistent with each accession having two *S*-alleles each derived from *P. fruticosa* and *P. avium* (Table 1, Fig. 2). The homoeolog dosage for the *S*-locus region in ‘Érdi Jubileum’ was 1A:3B, which was consistent with it sharing only one *S*-allele with *P. fruticosa* and three *S*-alleles with *P. avium.* (Table 1, Fig. 2).

**Table 1:** Overview of subgenome expression bias for homoeolog pairs for four comparisons: A vs. A’, A vs. B, A’ vs. B and AA’ vs. B. Values represent medians across all tissues and accessions, and the median log2FC values for each set. Homoeolog pairs with log2FC less that -3.5 or greater than 3.5 are identified by * and **, respectively.

|  | **Comparison (gene 1 / gene 2)** |  |  |  |
| --- | --- | --- | --- | --- |
|  | **A/A'** | **A/B** | **A'/B** | **AA'/B** |
| All pairs in analysis |  |  |  |  |
| Mean number | 5321 | 7050 | 6551 | 8084 |
| Median log2FC | -0.045 | -0.139 | -0.122 | -0.234 |
| All pairs biased towards gene 1* |  |  |  |  |
| Mean number | 139 | 103 | 99 | 94 |
| Median log2FC | -5.49 | -5.51 | -5.55 | -5.49 |
| All pairs biased towards gene 2** |  |  |  |  |
| Mean number | 126 | 154 | 158 | 3 |
| Median log2FC | 5.07 | 5.14 | 5.65 | 4.11 |

### Subgenome expression bias

Pairwise comparisons of the expression of homoeologs, normalized for dosage (see Methods), indicate that the expression of the A and A’ subgenomes is fairly balanced, while the A/B and A’/B comparisons show an overall expression bias towards the A and A’ subgenomes, respectively (Table 2, Supplementary Table S3, Supplementary Fig. S3 and S4). The overall median log2FC for A/B gene pair comparisons was -0.139, and the overall median log2FC for A’/B gene pair comparisons was -0.122 (Table 2, Supplementary Table S3). This bias was consistent across all five tissues and six accessions (Supplementary Fig. S3 and S4).

**Table 2:**
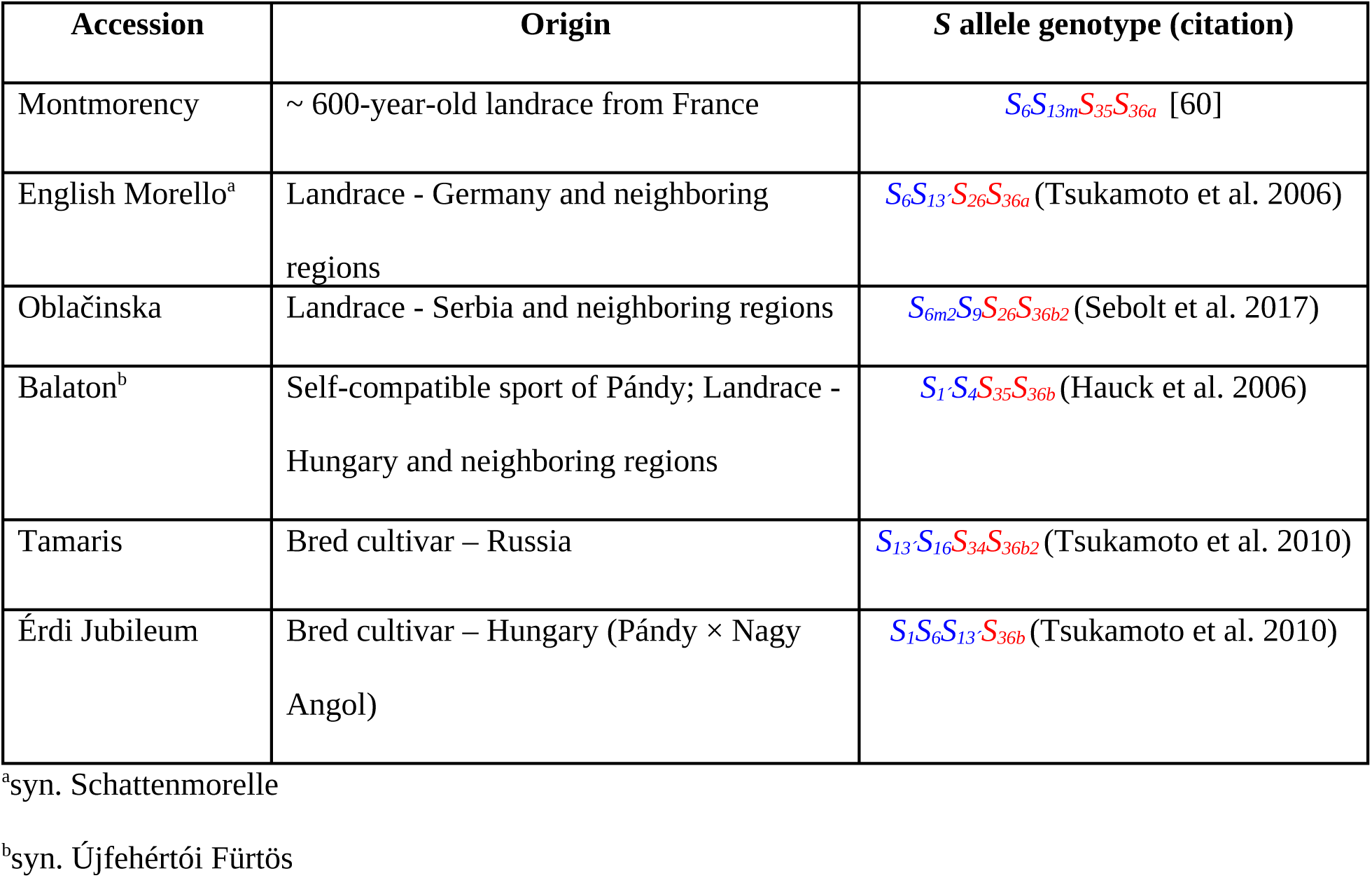
Sour cherry accessions included in this work, their origins and previously determined *S*-allele genotypes [35]. Five of the *S*-alleles present in the accessions (*S*_1_, *S*_4_, *S*_6_, *S*_9_, and *S*_13_) have only been identified previously in sweet cherry [58,59]. Goeckeritz et al. (2023) identified *S*_35_ and *S*_36a_ in *P. fruticosa*. *S*_26_ and *S*_34_ have only been identified in sour cherry, suggesting that they are derived from *P. fruticosa*.

When we treated the A and A’ subgenomes as one, still normalizing for dosage, the expression bias was strongly in favor of AA’, with an overall median log2FC of -0.234 (Table 2, Supplementary Table S3, Fig. 3). This AA’ bias was consistent across all tissues and accessions, with ‘Érdi Jubileum’ consistently exhibiting the highest expression bias (Supplementary Table S3, Fig. 3). Among the five tissue types evaluated, fruit stages 1 and 2 showed the most variation across the accessions for the relative magnitude of the AA’ bias (Fig. 3, Supplementary Table S3). Specifically, for fruit stage 1, the median log2FC ranged from -0.173 for ‘Balaton’ to -0.724 for ‘Érdi Jubileum’, and for fruit stage 2, the median log2FC ranged from -0.242 for ‘Montmorency’ to -0.731 for ‘Érdi Jubileum’ (Supplementary Table S3). Except for ‘Montmorency’, within individual accessions, the magnitude of AA’ homoeolog bias changed significantly over the course of fruit development, peaking at different development stages depending on accession (Fig. 3, Supplementary Table S3). For example, ‘Tamaris’ AA’/B gene pair comparisons had median log2FC values of -0.66 in fruit stage 1, -0.627 in fruit stage 2, and -0.22 in fruit stage 3. By comparison, ‘Balaton’ AA’/B gene pair comparisons had median log2FC values of -0.173 for stage 1, -0.682 for stage 2, and -0.22 for stage 3. However, it should be noted that all six accessions had similar AA’/B gene pair bias levels for fruit stage 3 (Supplementary Table S3) Non-reference level homoeolog pairs (i.e. dosage other than 1A:1A’:2B) resulting from homoeolog exchange or chromosome replacement detected in ‘English Morello’, ‘Tamaris’, and ‘Érdi Jubileum’ did not show a significant difference in subgenome expression bias (Supplementary Fig. S5).

**Figure 3:**
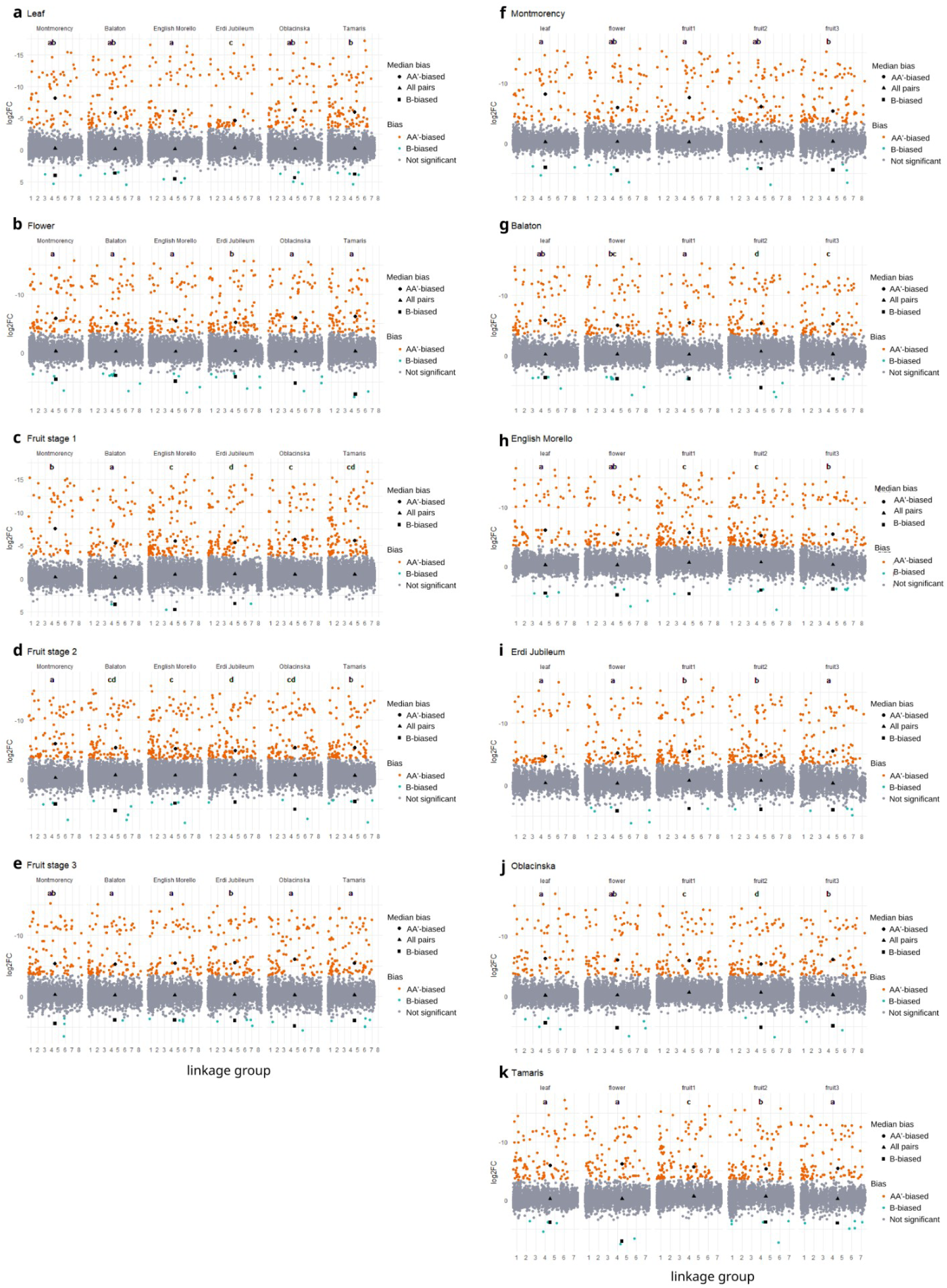
Dot plots of log2FC values for dosage-normalized AA’ versus B homoeolog pair expression comparisons for the five tissues (a-e) and six accessions (f-k). AA’ and B homoeolog pairs with a log2FC less than -3.5 or greater than 3.5, respectively, are colored according to the subgenome towards which they are biased. Median log2FC values for biased homoeolog groups and all homoeolog pairs are indicated in black. Significance groups based on analysis of variance and Tukey’s HSD of log2FC values are indicated by the letter above each dotplot, with “a” representing the least amount of expression bias and subsequent letters denoting higher levels of significance in expression bias. **Alt text:** A series of dot plots, labeled a-k. Each panel encompasses either one tissue for all accessions or one accession for all tissues. The x axis is chromosomes and the y axis is log2(Fold Change), and each dot is the log2(Fold Change) of one homoeolog pair. Statistical significance is noted above each dotplot by a letter.

The disparity between the number of significantly AA’-biased and B-biased homoeologs was immediately striking (Table 2, Fig. 3). Homoeolog pairs were considered significantly biased when the log2FC between homoeologs was greater than 3.5 or less than -3.5 based on Bird et al. (2021). A median of 87.5 homoeolog pairs were significantly biased towards the AA’ subgenomes across all accessions and tissues, while a median of 3 homoeolog pairs were significantly biased towards the B subgenome (Table 2, Supplementary Fig. S6, Supplementary Tables S4 and S5). Given the relatively small numbers of AA’-biased homoeolog pairs we then examined what fractions of the biased pairs were in common across the six accessions and found that relatively few of the AA’-biased homoeolog pairs were shared across accessions. The percentages of shared AA’-biased homoeolog pairs ranged from 7.4% (n=22) for fruit stage 2 to 14.6% (n=28) for flowers (Supplementary Fig. S6). In contrast, relatively more AA’-biased homoeolog pairs were unique to one accession, ranging from 33.2% (n=79) for leaves to 57% (n=170) for fruit stage (Supplementary Fig. S6).

### Subgenome-specific expression of four expansin genes expressed during fruit ripening

Using the ‘Montmorency’ reference genome, we identified four expansin genes previously identified in an RNA expression analysis of ‘Montmorency’ fruit ripening (*exp*1, 2, 4 and 5) [36]. For each gene, three homoeologs were identified which mapped to each of the three subgenomes. The *exp*5 and *exp*1 homoeologs mapped to chromosomes 1 and 2, respectively, while *exp*2 and *exp*4 were within 3Mb of each other on chromosome 6 (Fig. 2). The results of a coding sequence phylogeny of the ‘Montmorency’ expansin homoeologs with their corresponding orthologs from *P. avium* and *P. fruticosa* were consistent with their known ‘Montmorency’ subgenome placements (Fig. 4). Homoeologs on the ‘Montmorency’ A and A’ subgenomes clustered closest to their orthologs in *P. fruticosa* and homoeologs on the ‘Montmorency’ B subgenome clustered closest to their respective orthologs in *P. avium*. The one exception was the ‘Montmorency’ *exp*4 on subgenome A, which was equally distant from both progenitors. A closer examination of the coding sequence of the chromosome 6A *exp*4 shows it has several unique variants not shared with either progenitor, and therefore a putative progenitor origin could not be assigned for this homoeolog (Supplementary Fig. S7).

**Figure 4:**
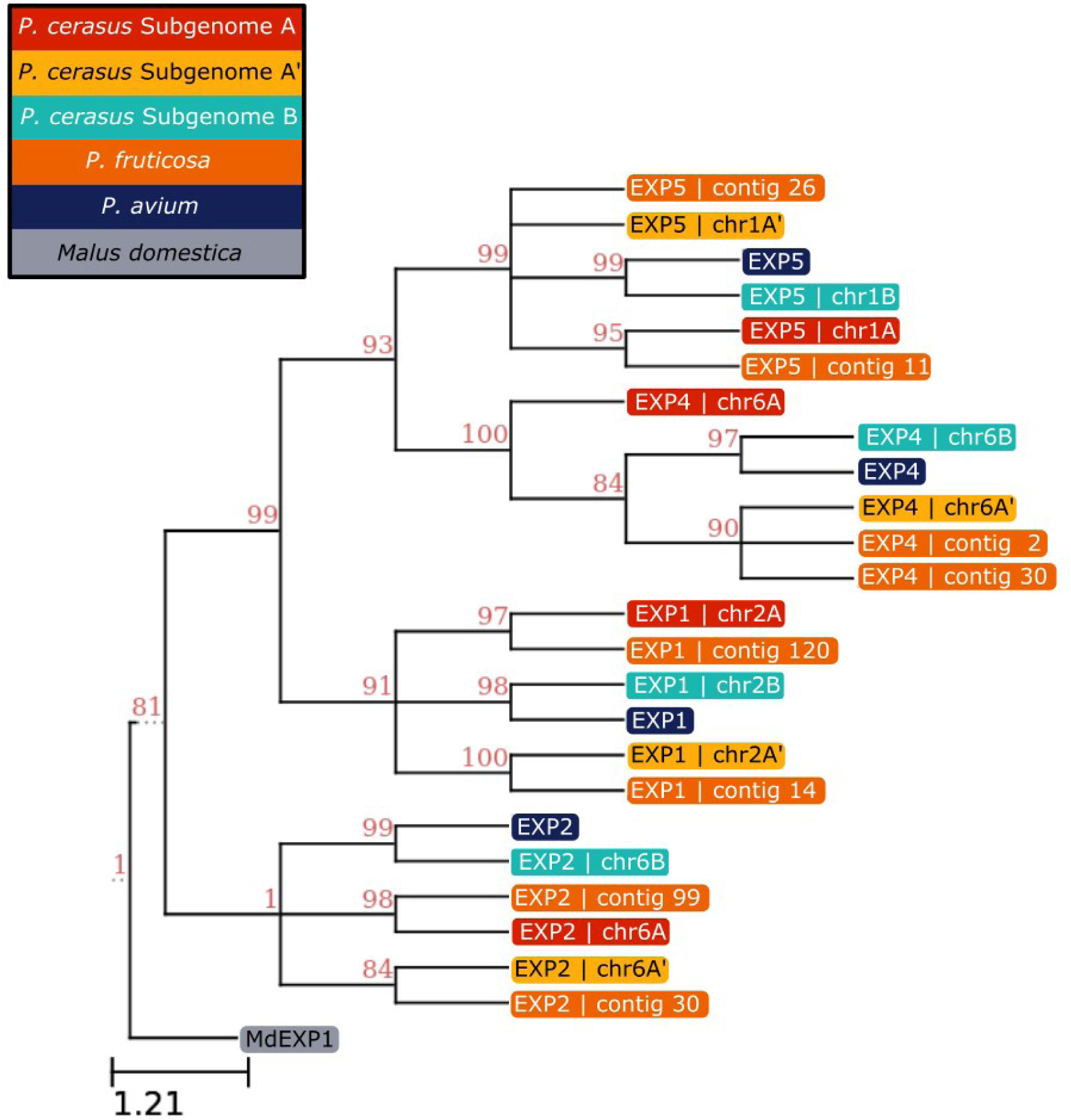
Phylogeny of the coding sequences for the homoeologs of four ‘Montmorency’ expansin genes first characterized in Yoo et al. (2003) and their corresponding orthologs in sweet cherry (Tieton v2.0 reference genome, Wang et al. 2020) and *P. fruticosa* (Goeckeritz et al., 2023). The subgenome location of each expansin homoeolog (A, A’ or B) is identified and color coded. An expansin from apple (*Malus × domestica*) is included as an outgroup (NCBI gene ID: 103447269). **Alt text:** A phylogenetic tree of expansins based on their coding sequences. Each expansin is color-coded according to its sour cherry subgenome or species.

All four ‘Montmorency’ expansin homoeologs were most highly expressed in fruit stage 3 compared to stages 1 and 2 (Supplementary Fig. S8). However, the *exp*4 homoeologs showed comparatively lower expression in stage 3 compared to that of the three other expansins. We attribute this to our fruit sampling time missing peak expression, as Yoo et al. (2003) showed *exp*4 expression declining at the very end of fruit development. As our RNAseq results were consistent with the qPCR findings of Yoo et al. (2003) and showed expansin expression mainly in the final stage of fruit development, we only used data from fruit in stage 3 in our analyses of subgenome expression bias.

Homoeolog expression for all four expansin genes was consistent with the subgenome presence-absence predictions illustrated in the karyotype analysis (Fig. 2, Fig. 5, Supplementary Fig. S9). To visualize this association between DNA presence/absence variation and gene expression, chromosome coverage for both the DNA and RNA short read alignments are shown in tandem for each of the homoeologs for each gene (Fig. 5 and Supplementary Fig. S9). As expected, homoeologs predicted to be missing due to chromosome replacements and/or HE show negligible expression, and the homoeologs with higher predicted dosage have higher expression. Without this knowledge of variation in homoeolog dosage, we might have falsely concluded that there was subgenome expression bias towards one of the homoeologs, with the assumption of each parental copy being present at similar ratios.

**Figure 5:**
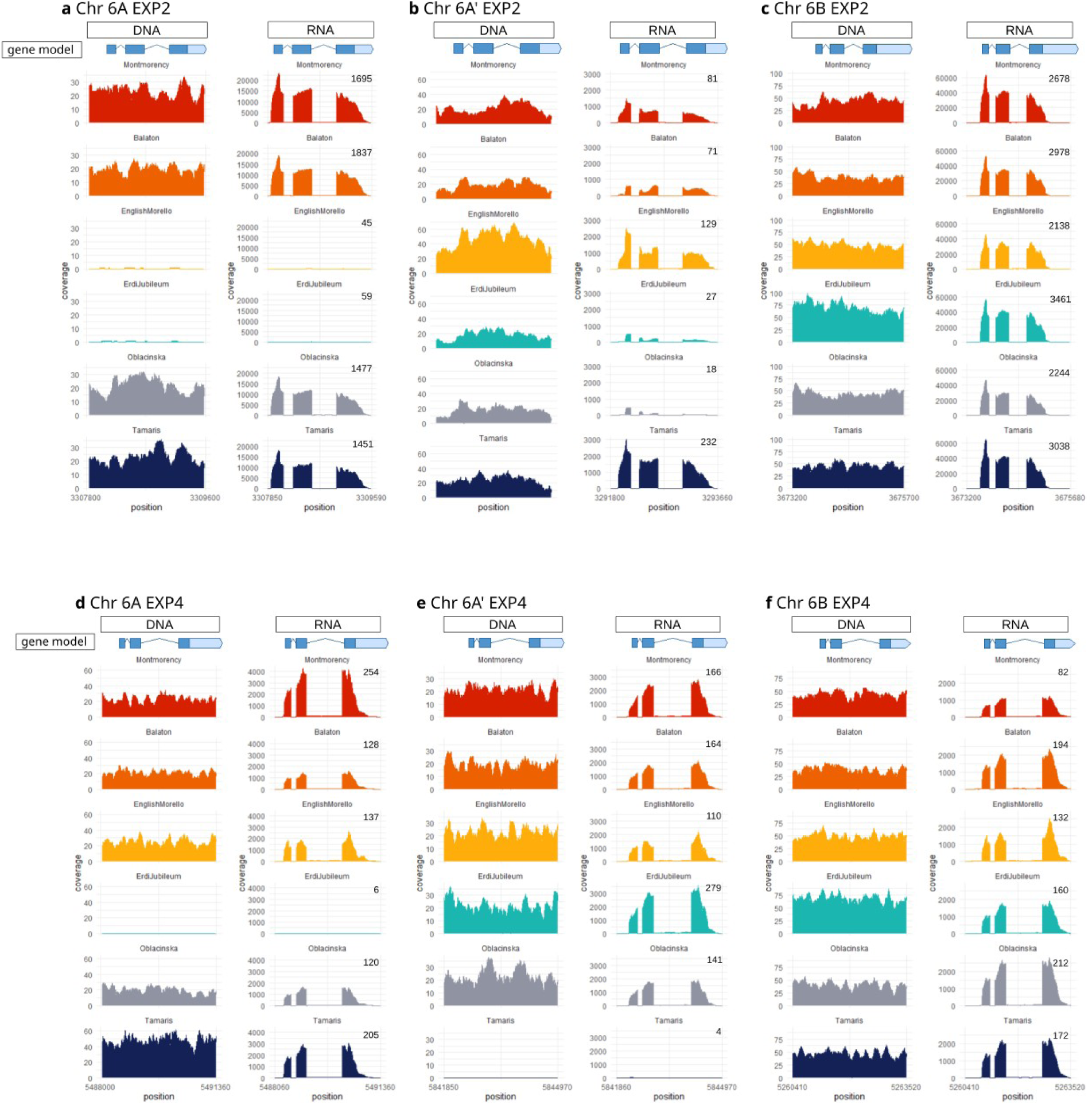
Alignment and coverage of DNA short reads and RNA-seq short reads for each sour cherry accession to the *exp*2 and *exp*4 gene regions (a-c and d-f, respectively) from the ‘Montmorency’ reference genome. RNA sequence data is from fruit in development stage 3. TPM is included on each RNA coverage graph. Gene IDs and TPM numbers are compiled in Supplementary Table S6. **Alt text:** A series of graphs showing DNA sequence and RNA sequence coverage for *exp2* and *exp4* on each homoeolog for each accession.

Of the four expansin genes investigated, only *exp*2 showed evidence of subgenome expression bias. In the accessions with 1:1:2 dosage of *exp*2 (i.e. ‘Montmorency’, ‘Oblačinska’, and ‘Balaton’), all exhibit significant bias against the A’ allele in the A/A’ and A’/B homoeolog comparisons in fruit stage 3 (Fig. 5, Supplementary Table S6). ‘Tamaris’ also exhibits differential expression between the A and A’ and A’ and B *exp*2 homoeologs, but it did not meet our log2FC threshold of +/-3.5 (A/A’ log2FC: -2.762; A’/B log2FC: 2.801). In the two accessions that have variation in subgenome *exp*2 dosage, ‘English Morello’ and ‘Érdi Jubileum’, the dose-normalized expression of the remaining homoeologs is also strongly biased against the A’ *exp*2 allele in favor of the B *exp*2 allele. This is despite having different *exp*2 homoeolog dosage for ‘English Morello’ and ‘Érdi Jubileum’ of 0A:2A’:2B and 0A:1A’:3B, respectively. We did not identify any variants within the DNA or mRNA sequence of the subgenome A’ *exp*2 or its promoter region that would explain lowered expression, suggesting the mechanism of suppression may be epigenetic or more distal to the locus. Further work is needed to determine if the observed dosage changes in *exp*2 homoeologs, i.e. the absence of the A-homoelog, increased expression of the B-homoeolog due to HEs, and reduced expression of the A’-homoeolog presumably due to expression bias, affect the phenotype. Interestingly, other genes that may play a role in fruit softening, including pectin methylesterase and pectinesterase inhibitors, also show an expression bias in favor of the B-homoeolog in fruit growth stage 3 (Supplementary Table 5).

## Discussion

### Genome-wide variation in subgenome dosage

The previously published finding that the allotetraploid sour cherry cultivar Montmorency is composed of three subgenomes was based on the finding that the *P. fruticosa*-like ancestor of ‘Montmorency’ was very likely an allopolyploid contributing two divergent subgenomes, A and A’ [32]. Here we present the subgenome structure for six additional sour cherry accessions, and the results for all six are consistent with our hypothesis that sour cherry consists of three subgenomes: A, A’ and B. The finding that a majority of the HEs identified were between the *P. fruticosa*-derived subgenomes versus exchanges between the *P. fruticosa-* and *P. avium-*derived subgenomes is consistent with prior work that reported preferential pairing of these subgenomes at meiosis [28]. For example, ‘Tamaris’ exhibited 11 exchanges between the A and A’ subgenomes and only one exchange between the A and B subgenomes. The resulting HE patterns in ‘English Morello’ and ‘Tamaris’ in particular suggest that over time, the two *P. fruticosa*-derived genomes in sour cherry may become a mosaic of A and A’ chromosomal regions. However, an examination of the centromeric regions suggests that the patterns may differ significantly between individuals. For example, the centromeric region identity for ‘English Morello’ chromosome 4 is 2A:2B while the centromeres for ‘Tamaris’ chromosome 4 are 2A’:2B.

The subgenome dosage pattern of the cultivar ‘Érdi Jubileum’ illustrates the extreme genome structure variation resulting from chromosome replacements that can occur when breeding for new selections. Notably, in four of the eight chromosome groups, a *P. fruticosa*-like chromosome was replaced by a *P. avium*-like chromosome, resulting in a 3× dosage of the *P. avium*-like subgenome (chromosomes 3, 6, 7 and 8). Concurrently, chromosome group 2 in ‘Érdi Jubileum’ has a 3× dosage of the *P. fruticosa* subgenomes. The 3× dosage of large portions of five of the eight chromosomes could be the result of irregular chromosome pairing at meiosis in one or more of the parental gametes, or the result of a *P. avium* introgression further back in its pedigree. From a breeding standpoint, the extreme deviation from the 1:1 *P. fruticosa* to *P. avium* subgenome ratio exhibited by this bred cultivar indicates that selection pressure from modern plant breeding can result in a dramatic increase in unbalanced allotetraploid subgenome structure.

The ‘Montmorency’ reference genome showed no evidence of HEs between the ancestral subgenomes based on the combined results from PacBio long reads, Hi-C sequencing used in assembly, and kmer analysis of the assembled subgenomes [32]. In our work, ‘Balaton’ and ‘Oblačinska’, both landraces, also showed no evidence of HEs when short-read DNA sequences were aligned to the ‘Montmorency’ reference. As the *P. fruticosa*-derived A and A’ subgenomes preferentially pair at meiosis, we would expect to see at a minimum, evidence for crossing over between these two subgenomes in successive generations following the initial interspecific hybridization event. The fact that we did not identify any A/A’ crossover events in three of the four landrace accessions suggests that either sour cherry may possess a genetic mechanism that limits homoeologous exchange [46], or the three landraces may be first-generation allotetraploids. The scenario where three separate landraces of sour cherry are all first-generation interspecific hybrids strikes us as improbable, therefore we also considered the possibility that we may have underestimated the number of HEs in ‘Balaton’ and ‘Oblačinska’. It is possible that, with only short-read sequencing, we may have failed to find HE events that would have been detected by high quality long read sequencing. All the HEs we reported were between 5 Mb and 25+ Mb in size, and it is possible that the five surveyed accessions could contain smaller HE events and/or events that are balanced between the subgenomes. These events, which would not result in dosage changes detectable on the coverage graphs [47], were not identified in this study using short-read data and our bioinformatic pipeline. In the only other sour cherry reference genome published to date, the German landrace ‘Schattenmorelle’, 28 HEs were identified [48]; however, all the HE regions were less than 5 Mb and therefore significantly less than the HEs we identified.

The unbalanced three subgenome structure identified in sour cherry [32] along with the prevalence of HEs and subgenome chromosome replacements provides an explanation for the low fertility and poor fruit set prevalent in sour cherry compared to its diploid sweet cherry progenitor. Studies of meiosis in sour cherry show varying rates of univalents and multivalent chromosome pairing, resulting in aneuploid and unbalanced gametes – suggesting sour cherry has not reached cytological diploidization [19,20,22,27]. Additionally, sour cherry exhibits disomic and tetrasomic patterns of inheritance, further supporting occasional pairing between homoeologs at meiosis [28,34]. Together, these findings are consistent with prior speculation that sour cherry is a neopolyploid where selection for increased fertility has yet to be achieved [32]. Unfortunately, the consequences of the low gamete fertility in cherry are particularly severe as cherry is not a multi-seeded fruit such as tomato or blueberry. Instead, cherry and all *Prunus* species have just two ovules, one of which typically degenerates, leaving one seed inside the endocarp. Fruit set requires successful fertilization and the initiation of embryo development. Therefore, any disruption that results in a non-viable ovule or zygote will result in the flower failing to develop into a fruit [49].

The evolution and reproductive biology of sour cherry may offer clues as to why the allotetraploid sour cherry has failed to achieve proper bivalent pairing in meiosis. Firstly, sour cherry likely formed from an unreduced (2*n*) gamete from sweet cherry and a reduced gamete (*n*) from an allotetraploid ground cherry [23]. It may be more difficult to achieve diploidization with an initial 1:1:2 subgenome structure as opposed to an initial whole genome duplication with subgenomes present in equal proportions. Most genomically-characterized allopolyploids are thought to be the result of a whole-genome duplication event that occurred before or after, not simultaneous with, the hybridization event. Secondly, the long life cycle in sour cherry plus its ability to spread through vegetative suckers would have enabled the newly-formed interspecific hybrids to persist and spread despite their low fertility.. Finally, sour cherry has been shown to cross with its progenitor species where their ranges overlap [50], and this continued introgression would have likely disrupted the movement towards diploidization.

As our study focused on sour cherry landraces and cultivars, it is important to consider the impact that human selection may have had on their subgenome structure. All four landraces are known to have been vegetatively propagated for hundreds of years, and represent clonal lineages that express subtle phenotypic differences likely due to the accumulation of epigenetic changes and somatic mutations. This raises the possibility that individuals with a 1A:1A’:2B subgenome ratio would exhibit relatively high fertility compared to subsequent offspring. To date, modern breeding has failed to develop any sour cherry cultivars that have the high fruit set and thus yield of ‘Montmorency’. Despite the low fertility in sour cherry breeding germplasm, one of the advanced generation hybrid populations in the MSU sour cherry breeding program has individuals with exceptionally high fruit set. The ability to obtain DNA sequence and characterize the genome structure in sour cherry finally provides breeders with the tools to explore the underlying basis of fertility differences in sour cherry germplasm.

### Subgenome expression bias

Subgenome expression bias across all six sour cherry accessions and tissues was consistently in favor of the A and A’ subgenomes compared to the B subgenome. This consistent presence and direction of subgenome expression bias across accessions is similar to results in other species that show expression bias favoring one subgenome establishing immediately after polyploidization and persisting through generations [17,51]. We initially looked at 1A:1B and 1A’:1B subgenome expression comparisons and observed consistent bias favoring gene expression of the A or A’ subgenome over the B subgenome at magnitudes similar to other subgenome expression bias publications [51]. However, we found that treating the two subgenomes inherited from the tetraploid ancestor (A and A’) as one unit and comparing them with the subgenome inherited from the diploid ancestor (B) nearly doubled the magnitude of the subgenome expression bias.

We speculate that the consistent subgenome bias for the combined A+A’ *P. fruticosa* subgenomes over the *P. avium* subgenome may reflect a legacy of expression optimization that evolved in the allotetraploid *P. fruticosa* and not in the diploid sweet cherry. Specifically, unlike the B subgenome, the A/A’ subgenomes would have already adjusted to a 4× nucleus. Models indicate that subgenome expression bias is in fact the expected consequence of a neoallopolyploidy event, especially when parental subgenomes are of different sizes [52], and tetraploid *P. fruticosa* and diploid *P. avium* are ∼660Mb and ∼330Mb, respectively. Interestingly, in newly synthesized hexaploid wheat, gene expression was strongly biased in favor of the A and B subgenomes when they were treated as a unit, as we have done with the A and A’ subgenomes [53]. The authors hypothesize that the strong AB dominance over the D subgenome in newly-synthesized polyploid lines may be an effort to balance gene expression, as the parental DD line exhibited much higher gene expression than the parental AABB line.

For all six accessions and tissues examined, we found extensive variation for which genes exhibited significant expression bias. We attribute at least part of this lack of overlap to sampling variation, as the plants were grown in the field and logistics required that they be sampled sequentially rather than simultaneously. However, the variation in which genes exhibited expression bias could also reflect genetic differences among the accessions as they were chosen to represent a range of diversity. Additionally, it is possible that part of the variation could be due to differences in each accession’s manifestation of the ‘genetic shock’ that has been shown to accompany allopolyploidization followed by differing selection pressures. These different selection pressures could have an environmental basis as the accessions used in this study originated from a wide range of environments, i.e., the relatively warmer Mediterranean region, north to Germany and east into Russia. *P. fruticosa* is significantly more cold hardy and later blooming than *P. avium* and therefore more adapted to the cold Russian environment where it evolved. The two species also exhibit a suite of morphological traits that differ dramatically, including leaf size, shape, and gland characteristics. A morphological analysis of sour cherry germplasm grown in a common orchard in Michigan (the same location as the accessions in this study) found a greater morphological resemblance to *P. fruticosa* for germplasm that originated in the colder range for sour cherry and vice versa for germplasm collected in the milder sour cherry range [54]. Selective forces may have contributed to the patterns of morphological variation as well as the differences in biased homoeologs in our study.

### Differences in subgenome structure variation for four expansion genes

Our finding that subgenome dosage of expansin genes frequently diverges from 1:1:2 illustrates why knowledge of subgenome dosage is critical to making breeding decisions such as designing crosses and implementing selection strategies. Additionally, gene expression predictions would be optimized with an accurate assessment of subgenome bias. For example, the A-subgenome homoeolog of *exp*2 is absent in both ‘English Morello’ and ‘Érdi Jubileum’, and if homoeolog expression was assumed to be consistent with a 1A:1A’:2B subgenome ratio, a conclusion of subgenome bias against the A’ *exp*2 homoeolog in fruit stage 3 would have been false. The consistent subgenome bias in favor of the *P. fruticosa* subgenomes also could be leveraged to optimize breeding outcomes. For example, the introgression of desirable genes into the *P. fruticosa* subgenome as opposed to the *P. avium* subgenome may result in higher expression of these introgressed genes. Alternatively if a desirable gene were introgressed into the *P. avium* subgenome it may be more likely to have suppressed expression.

Our study was limited in that we did not functionally validate the biological consequences of the four expansin genes, namely their potential role in fruit softening during the final stage of fruit ripening. However, as increased fruit firmness is an important target for both sweet and sour cherry breeding, our findings suggest that further investigations of whether any of these four expansins play a role in fruit softening would be useful. In addition, it is interesting to consider fruit firmness in cherry in the larger context as fruit firmness differs dramatically between species, and along with fruit size, it is a major trait associated with domestication [55,56]. The fruit of wild sweet cherry (also *P. avium*) is soft and small while all domesticated sweet cherries have fruit that is firmer and larger than the wild ancestor. Like wild *P. avium*, *P. fruticosa* has soft fruit and we have no knowledge of any firm fruited *P. fruticosa* selections. All sour cherries have fruit that is softer than commercially grown sweet cherries. The softer fruit and associated juiciness of sour cherry make it well suited for processed products such as juice, jams and pies, while the firmer sweet cherry is well suited for the fresh market. There are subtle differences in fruit firmness among sour cherry selections; however, no sour cherry selections have firmness levels approaching sweet cherry. Of the sour cherry cultivars available, ‘Érdi Jubileum’, bred in Hungary for the fresh market, is one of the most firm-fruited sour cherry cultivars available [57]. It is intriguing to consider that the increased *P. avium* subgenome contribution in ‘Érdi Jubileum’ may be associated with this increased firmness, either through the addition of more desirable sweet cherry alleles or as a strategy to counter the prevalent expression bias in favor of alleles on the *P. fruticosa*-derived subgenomes.

## Conclusion

The subgenome structure for sour cherry accessions presented herein suggests that sour cherry is a neopolyploid that is still meiotically unstable. The increased number of HE and chromosome replacements in modern bred cultivars suggest that breeding may inadvertently act to further destabilize the subgenome structure. In addition, we documented a consistent direction of subgenome expression bias across all accessions and tissues that supports the growing hypothesis that if subgenome expression bias exists, it tends to act in a consistent direction across individuals within a species [17,51,53]. These results have implications both for the evolutionary genomics of polyploidy and for sour cherry breeders seeking to better characterize their germplasm to inform breeding decisions.

## Materials and Methods

### Plant materials and tissue collection

The six plant materials examined include four landrace accessions: ‘Montmorency’, ‘Balaton’, ‘English Morello’, and ‘Oblačinska’; and two cultivar accessions: ‘Tamaris’ and ‘Érdi Jubileum,’ bred in Russia and Hungary, respectively (Table 1). All accessions were grown at the Michigan State University Clarksville Research Center in Clarksville, Mich. All tissues collected (Supplementary Fig. S1) were flash-frozen in liquid nitrogen and stored at -80℃ until extraction.

Young leaves were collected for DNA extraction. To minimize the effects of circadian rhythm on our subgenome expression bias analyses, we collected tissues for RNA extraction within the same three hour window each collection day. Three biological replicates of the following tissues were collected for RNA extraction: young leaves, whole flowers at the “balloon” stage, and developing fruit collected weekly starting at anthesis. Once the fruit were collected, the widest diameter of the fruit was measured and used to create a fruit growth curve that allowed us to match similar developmental stages across accessions despite the differences in the timing of fruit development and ripening. Based on the fruit growth curves (Supplementary Fig. S1), three stages of fruit development were selected for RNA sequencing for each accession. The stages selected were the start of the first exponential growth phase (stage 1), the end of the growth-plateau phase of pit hardening (stage 2), and the peak of the second exponential growth phase concurrent with fruit ripening (stage 3).

### DNA and RNA extraction and sequencing

DNA was extracted using the Qiagen Plant DNeasy kit (www.qiagen.com) according to manufacturer’s instructions. Libraries were prepared by the MSU Research Technology Support Facility (RTSF) genomics core using the Illumina TruSeq Nano DNA Library prep kit according to manufacturer’s instructions (www.illumina.com). Short-read sequencing (paired-end, 150bp) was also performed at the MSU RTSF genomics core on an Illumina HiSeq4000 to a depth of ∼40× per sample. RNA was extracted using a CTAB-based method [61]. RNA libraries were prepared with the Illumina TruSeq Total mRNA kit according to manufacturer’s instructions and sequenced 2×150bp on a HiSeq 4000 at the MSU RTSF to a target of ∼35 million reads per library.

### Assigning subgenome dosage

Illumina 2×150bp DNA reads for each accession were trimmed for quality using Trimmomatic v0.38 [62] and aligned to the 24 scaffolded chromosomes of the ‘Montmorency’ reference genome [32] using BWA mem v0.7.17 on default settings [63]. The resulting sequence alignment file was sorted with SAMtools v1.15 [64] and PCR duplicates were marked with GATK v4.0 markduplicatesspark [65]. SAMtools depth was then used to call sequence depth values for each base pair of the ‘Montmorency’ reference where the aligned read base quality and mapping quality were both greater than 20. These depth values were then averaged over 1 kb windows on each chromosome to obtain average sequence depth. Sites of HE were assigned based on reciprocal changes in sequence coverage between homoeologs. Sequence depth values averaged over 1 Mb windows for each chromosome for each accession were graphed in R using package ggplot2 [66,67]. The resulting subgenome karyotypes for the six accessions were visualized using Inkscape v1.0.1 (https://inkscape.org).

### Gene expression counts and dose normalization to compare the A/A’, A/B, and A’/B subgenomes

RNA sequence reads were aligned to the 24 chromosome ‘Montmorency’ genome using STAR v2.7.9a [68] and quantified using StringTie v2.1.3 [69] on expression estimation mode with multi-mapping correction. Stringtie abundance files for each sample were imported into R v4.2.2 [67] and combined with the homoeolog dose/pair assignment files using tidyverse v2.0.0 packages [70]. All homoeolog pairs with a combined transcripts per million (TPM) less than 10 were discarded. Gene expression counts for all three tissue replicates for each tissue type were combined, only including homoeolog pairs found to be expressed in all three replicates, and average TPM for each gene was calculated. We then divided the average TPM for each gene by its dosage to obtain the dose-normalized gene expression value which was used for homoeolog expression comparisons.

### Subgenome expression bias in 1:1 subgenome comparisons (A/A’, A’/B, A/B)

Using the dose-normalized average TPM values, log2FC was calculated as log_2_(TPM-gene2+0.01/TPM-gene1+0.01). In all results a negative log2FC value indicates that the first homoeolog in the pair is more highly expressed, and a positive log2FC value indicates that the second homoeolog in the pair is more highly expressed. Comparisons were done for three sets of homoeolog pairs: A/A’, A/B, and A’/B.

### Subgenome expression bias comparison between AA’ and B homoeolog sets

To compare expression between both *P. fruticosa*-derived homoeologs and the *P. avium*-derived homoeologs, we selected gene pairs that were present in the A/A’ homoeolog pair set and also present in the A/B homoeolog pair set. As described above, only homoeolog pairs that were expressed in all three tissue replicates were included. In cases where the homoeolog genotype was AA’BB, the TPM sum for the A + A’ homoelogs and the TPM for B homoeologs were both dose-normalized by dividing by 2. In cases where the homoeolog genotype was ABBB or A’BBB, the sum for the A + A’ homoeologs was dose-normalized by dividing by 1 and the TPM for the B homoeolog was dose-normalized by dividing by 3. We then calculated the log2FC between the combined A/A’ homoeologs’ TPM and the B homoeolog TPM as log2FC (TPM geneB/(TPM geneA + TPM geneA’)).

### Statistical testing of subgenome expression bias comparisons and comparison among accessions

Analysis of variance on log2FC values were initially performed individually for each accession comparing all tissues and secondly for each tissue type comparing all accessions. Tukey’s HSD was used to assign significance groups to each tissue or each accession (p <0.05). Sets of homoeolog pairs with log2FC < -3.5 (AA’-biased) or log2FC > 3.5 (B-biased) for each tissue in each accession were compared across accessions using tools in the tidyverse R package [70].

Upset plots were created with R package UpSetR [71].

### Expansin location on chromosomes and phylogeny to determine progenitor relationships

We identified the four expansins (*exp*1, 2, 4 and 5) that were previously associated with ‘Montmorency’ fruit development [36] in the ‘Montmorency’ reference genome v1.0 [32] with a BLAST+ v2.7.1 search of the cds sequences [72]. Coding sequences for each expansin gene were also obtained from the *P. avium* cv. Tieton v2.0 reference genome [73] and the *P. fruticosa* reference genome [32] as independent representations for each progenitor. An ortholog from apple (*Malus* x *domestica*; NCBI ID: 103447269) was used as an outgroup, and all sequences were aligned with MUSCLE v3.8.31 [74]. Phylogenies were constructed with RAxML-NG v1.0 [75] using the GTR+G algorithm and 500 bootstrap replicates to create a consensus tree with all branches supported in at least 80 percent of replicates. Tree figures were generated using the ETE Tree Viewer (etetoolkit.org). *Exp*4 amino acid and coding sequence alignments were completed with MUSCLE v3.8.31 and figures were generated using R package ggmsa [76].

Transcript abundances for the expansins were taken from the whole-transcriptome Stringtie abundance datasets generated above. Using R v4.2.2 [67], we performed analysis of variance of expansin expression between subgenomes within each tissue and used a Tukey’s HSD test (p < 0.05) to determine statistical significance groups for expansin transcript abundances.

## Acknowledgements and Funding

We would like to thank Dr. Dave Douches and Dr. Courtney Hollender for their guidance on this project. We would also like to thank Audrey Sebolt, Dr. Brent Crain, Abby Seeger, and Jonas Berenkowski for their invaluable assistance in the field and lab. Dr. Joseph Hill and Dr. Chris Gottschalk also kindly provided support in the lab troubleshooting RNA extractions. This work was supported by the United States Department of Agriculture National Institute of Food and Agriculture (USDA-NIFA) project 2014-51181-22378.

## Author contributions

K.R. and A.I. developed the research questions and plan together. K.R., C.Z.G., A.Y., K.A.B. and P.E. jointly determined the methods. K.R. performed the analyses. K.R. wrote the first draft of the manuscript with input from A.I. and the final manuscript incorporated input from A.I., P.E., K.A.B, and C.Z.G. Editing was done by A.I. and K.R.

## Data availability

All DNA and RNA sequence data is uploaded to NCBI under project number PRJNA1251028 Bash and R scripts for analysis can be found at: https://github.com/KEBRhoades/pcerasus-subgenome-dosage-and-bias/

## Supplemental material available at Horticulture Research online

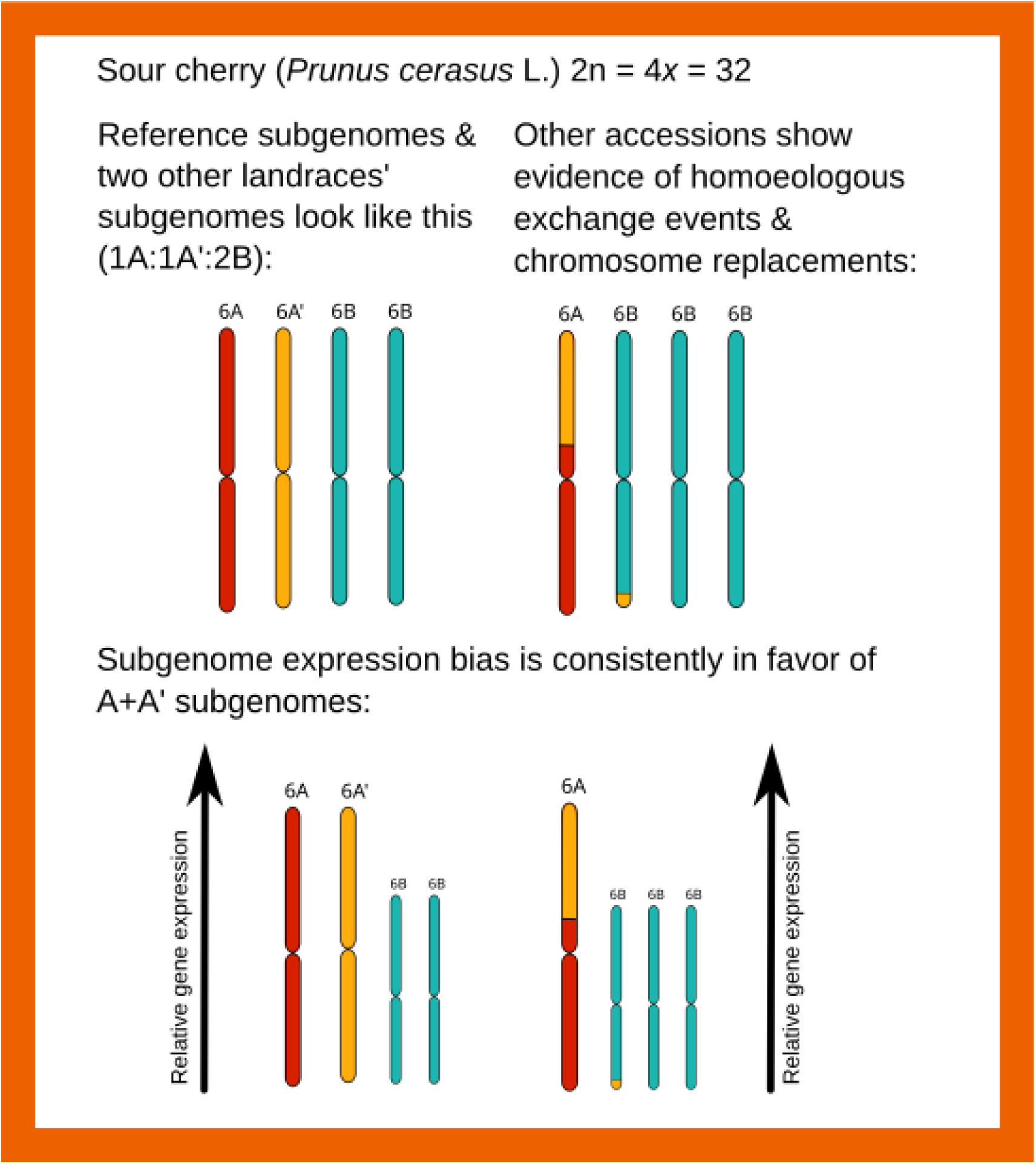

